# Categorical bias as a crucial parameter in visual working memory: the effect of memory load and retention interval

**DOI:** 10.1101/2022.02.23.481291

**Authors:** Cherie Zhou, Monicque M. Lorist, Sebastiaan Mathôt

## Abstract

Visual information can be stored as continuous as well as categorical representations in visual working memory (VWM) to guide subsequent behavior. Yet it is still unclear what determines whether VWM is represented as continuous or categorical information, or as a mix of both. Recent studies have shown that color VWM representations adjust flexibly depending on the number of memory items as well as the duration that these items need to be maintained for. The current study aims to extend and replicate these crucial effects. In a delayed estimation task, participants memorized one to four colored objects presented at different spatial locations, followed by a delay of 100, 500, 1,000, or 2,000 ms. Next, a probe indicated the location of the color that participants needed to report. We measured the extent to which responses were biased in the direction of prototypical colors. Crucially, we implemented this categorical bias in an extension to the classic mixture model (Zhang & Luck 2008) in which the center of the error distribution is a crucial parameter that characterizes the extent to which VWM is biased by color categories. We found that VWM shows a strong categorical bias in all cases, and that this bias increases with increasing memory load; strikingly, this effect of memory load on categorical bias is stronger at longer intervals (1,000 ms or longer), as compared to shorter intervals, yet it peaks for intermediate memory loads as opposed to the highest memory load. Overall, our results suggest that when visual information needs to be maintained for one second or longer, VWM becomes more reliant on categorical representations as memory load increases.

---

Visual working memory (VWM) stores relevant visual information to guide subsequent behavior. This information can be stored as a visual representation, in which case specific visual details, such as the exact shape, color, and orientation of a stimulus are retained; information can also be stored as a categorical representation, in which case a categorical tag, such as *a red coffee cup* is retained (Bae et al., 2015; Hardman et al., 2017; Olsson & Poom 2005). This distinction is related to a current debate about which brain areas are involved in VWM. According to some authors (Gayet et al. 2017; Harrison and Tong 2009), VWM mainly relies on sensory brain areas, which are presumably the areas that underlie visual representations (but see Ester et al., 2020); according to others (Bettencourt & Xu, 2016; Goldman-Rakic, 1995), VWM mainly relies on non-sensory, parietal and frontal, brain areas, which have been proposed to underlie categorical representations (Freedman et al. 2001; Lee et al.,2013; but see Ester et al., 2015). It is also conceivable that a mix of visual and categorical representations (Christophel et al. 2017; Huttenlocher et al., 2000), and thus a mix of sensory and non-sensory brain areas, is involved in VWM. However, if this is true, then a crucial and largely open question is which factors determine the relative contribution of visual and categorical representations in VWM maintenance.

The sensory-recruitment hypothesis posits that VWM relies on early visual (sensory) cortices: the areas that are involved in the initial encoding of visual input (Pasternak & Greenlee, 2005; Postle, 2006). In support of this view, studies using fMRI decoding have demonstrated that visual representations maintained in VWM can be decoded from activity in visual cortex over a delay period during which participants maintain information in VWM (Gayet et al., 2017; Harrison & Tong, 2009; but see Ritchie et al., 2017 for criticism on this method); this finding clearly shows that VWM information is represented, at least in part, in visual cortex. Studies using eye tracking have found indirect support for the sensory-recruitment hypothesis by showing that the contents of VWM affect very fast saccadic eye movements, which are believed to be driven largely by visual input (e.g., Theeuwes et al., 1998). For example, in a study by Hollingworth and colleagues (2013), participants maintained a color in memory, while executing a saccade to a target. The authors manipulated the color of the elements in the display such that either the target or a distractor matched the memory color. Importantly, the authors found that saccade landing positions were biased towards the memory-matching objects, regardless of whether they were targets or distractors; this bias occurred even for the earliest “express” saccades, suggesting that the contents of VWM modulate visual input at an early stage. Taken together, there is compelling evidence from neuroimaging (e.g., Gayet et al., 2017; Harrison & Tong, 2009) as well as eye-tracking (Hollingworth et al., 2013) that VWM relies on activity in early visual cortex.

In apparent conflict with the sensory-recruitment hypothesis, findings from some studies suggest that higher-order, frontal and parietal areas, instead of early visual areas, underlie the maintenance of VWM (Ester et al., 2015; Goldman-Rakic, 1995; Postle, 2006). Proponents of this view argue that, even though VWM representations can be stored in early sensory areas, these representations are susceptible to interference and not strictly necessary for VWM maintenance (Bettencourt & Xu, 2016). For example, Bettencourt and Xu (2016) found that, when distractors are expected during a retention interval, VWM-decoding performance in early visual cortex drops to chance level, while decoding performance in parietal cortex remains unaffected, as does behavioral performance. These results have been used to highlight the role of parietal cortex, instead of early sensory cortices, in VWM maintenance.

So far, studies seeking evidence for both hypotheses have mainly focused on continuous representations: representations that reflect the sensory properties of the stimuli (e.g., an exact hue of a color), rather than the categorical labels that are associated with them (e.g., ‘red’). Yet it is increasingly recognized that visual information can also be represented categorically in VWM (Bae et al., 2015; Hardman et al., 2017; Olsson & Poom, 2005; Panichello et al., 2019; Pratte et al., 2017; Souza & Skora, 2017). In a recent study, Bae and Luck (2019) asked participants to maintain an orientation in VWM, while they performed a visual discrimination task during the delay period, not unlike the paradigm by Bettencourt and Xu (2016). Importantly, Bae and Luck (2019) found that VWM became less precise and more biased by cardinal orientations when it was interrupted by the discrimination task. This suggests that the discrimination task interfered with the continuous component of the VWM representation, while the categorical component (i.e. the cardinal bias) was left intact. Phrased differently, continuous VWM representations are likely fragile and severely limited in capacity (e.g., Olivers et al., 2011), whereas categorical VWM representations are likely more robust and less limited in capacity. As mentioned earlier, a common view is that categorical information is represented in non-sensory brain areas such as parietal and frontal cortex (Freedman et al., 2001; Lee et al., 2013), which are associated with experience and learning of specific categories, whereas continuous information is represented in sensory brain areas such as early visual cortex. However, there may be considerable flexibility in the way in which VWM is represented in all brain areas, possibly depending on task demands (see Christophel et al., 2017 for a review). For example, recent studies have found evidence for continuous VWM representations also in frontal cortex (Ester et al., 2015), and, conversely, have found evidence for categorical VWM representations also in early visual cortex (Ester et al., 2020).

Studies on VWM, especially those using color, often use the “mixture model” that was introduced by Zhang and Luck (2008) to quantify performance on VWM tasks. This mixture model consists of a uniform distribution that represents random guesses, which are equally likely to come from any part of the color circle, and a normal (von Mises) distribution that represents “noisy” responses, which are centered on the memorized color but with random error (Wilken & Ma, 2004). This model can be used to estimate whether errors in VWM tasks are due to participants either not storing items in (or failing to retrieve items from) VWM (an increased influence of the uniform distribution), or due to decreased precision of VWM (a widening of the normal distribution).

Using variations of this method, several studies recently investigated the influence of continuous and categorical representations on VWM. For example, Hardman and colleagues (2017) asked participants to perform a delayed estimation task (Wilken & Ma, 2004), in which they first memorized one or more color stimuli; next, after a delay period, they reported the memorized stimulus on a color wheel. The authors added a new parameter to the mixture model that reflected the probability that items stored in VWM are continuous or categorical. They found that items became more likely to be represented categorically as memory load increased (see also Panichello et al., 2019; Pratte et al., 2017). Crucially, Hardman and colleagues (2017) found that a model in which memory representations were either categorical or continuous performed better than a related model in which memory representations were a weighted mix of both.

A related study by Bae and colleagues (2015) again used a delayed estimation paradigm to investigate categorical representations with or without a delay between the presentation of the memory display and the response wheel. Importantly, they found that memory responses were biased toward category centers even on no-delay trials, although this bias was bigger with a delay. From this, the authors concluded that representations are already categorically biased at the early stage of processing. This assumption has been further tested using neuroimaging and EEG-decoding techniques. For example, in an EEG study, Ester and colleagues (2020) generated time-resolved representations of the orientations that participants needed to categorize. They found that these representations were biased toward category centers as early as 300ms after stimulus onset. Consistent with this finding, Bae (2021) decoded EEG signals in a delayed location-estimation task. Crucially, the author found that decoded locations were categorically biased already during the perceptual encoding period and that this bias persisted into the retention period.

Here we tested the influence of categorical representations in VWM by implementing a parameter that represents the reliance on categorical representations in a straightforward extension to the classic VWM mixture model. We predict that, similar to the computational models of VWM from previous work (e.g., Hardman et al., 2017; Panichello et al., 2019; Pratte et al., 2017), this biased mixture model will better account for human performance on color-VWM tasks than the original mixture model does. We further predict that the reliance on categorical representations increases as a function of both the number of memory items and the duration for which these items need to be maintained. Specifically, we predict that when memory load is low, color VWM relies largely on continuous representations; however, because VWM has a limited capacity, especially for continuous representations, we predict that as memory load increases, VWM representations become more categorical. Moreover, we predict that shortly after the presentation of a stimulus, color VWM relies largely on continuous representations; however, because continuous representations are likely fragile and can be lost from VWM as the retention interval increases (Ricker & Cowan, 2010), we predict that VWM becomes progressively more categorical when visual information is maintained for a longer duration. We assume that categorical biases are a crucial component of VWM, and that it is important to establish these key effects in a high-powered study.

To test our predictions, we used a delayed estimation paradigm with different set sizes and delay periods. Participants were instructed to memorize the exact shade of one to four colors, followed by a delay of 100, 500, 1,000, or 2,000 ms. Next, an arrow appeared to indicate the location of the color that they needed to report. After the experiment, we asked participants to indicate their subjective color boundaries and prototypical colors (i.e. the color center) for different color categories; these values were used to analyze the results for each participant. We measured the *categorical bias*, which is defined as the response error coded such that positive values reflect a bias in the direction of the prototypical color of the category; a stronger categorical bias reflects a stronger reliance on discrete categories in VWM. Importantly, we implemented an extension of the mixture model (Zhang & Luck, 2008) that characterizes, for each participant separately, the extent to which VWM relies on categorical representations (see also Bae & Luck, 2019, who used a similar approach to measure biases away from cardinal orientations in VWM). Here we conceptualize categorical representation as an aspect of visual processing, which can occur without verbal labeling (Souza & Skora, 2017). We encourage nonverbal representations by instructing participants to memorize the *exact* colors and by providing feedback on their accuracy after each trial.

Traditionally, the two main opposing models of VWM are the slot model (e.g., Pratte et al., 2017; Zhang & Luck, 2008) and the resource model (e.g., Bays & Husain, 2008; Bays et al., 2009). The slot model posits that the capacity of VWM is limited to a fixed number of items; consequently, items would either be remembered or forgotten. In contrast, the resource model posits that VWM is a limited resource that can be flexibly distributed across as many items as need to be remembered; consequently, items would be represented with varying degrees of fidelity. Although it is not our primary aim to distinguish between these two models, our proposal to consider categorical bias as a crucial property of VWM combines elements of both: a purely continuous VWM representation without any categorical bias may function according to the resource model, in the sense that such representations can vary in their precision; in contrast, a purely categorical representation may function according to the slot model, in the sense that it is either remembered or not. In other words, slot and resource models may accurately describe different aspects of VWM. This idea is in line with the emerging view that VWM is not a unitary process, but rather consists of different types of representations (e.g., Christophel et al., 2017; Olivers et al., 2011). This idea is also consistent with the recent finding that a complex computational model that combines characteristics of both slot and resource models (i.e. a limited number of slots, each with a variable precision) outperforms traditional models of either kind, at least when considering performance on an orientation-memory task (Pratte et al., 2017). In summary, research on VWM is moving away from a strict theoretical dichotomy between slot and resources models, and is increasingly recognizing the importance of stimulus-specific biases (e.g., Christophel et al., 2017; Pratte et al., 2017).

## Hypotheses

We tested two main hypotheses using different memory loads and retention intervals:

1. The extent to which visual information is represented categorically in color VWM increases with increased memory load. Specifically, we hypothesized that when memory load is low, color VWM mainly relies on continuous representations, and will not, or hardly, be affected by categorical information; however, because of the limited storage capacity of continuous representations in VWM (Olivers et al., 2011; Zhang & Luck, 2008), we hypothesized that color VWM becomes progressively more categorical as memory load increases.
2. The extent to which visual information is represented categorically in color VWM increases with increased duration of the retention interval. Specifically, we hypothesized that when the retention interval is short, color VWM is not strongly affected by categorical information; however, as information is maintained for longer in VWM, we hypothesized that color VWM becomes progressively more categorical.

## Method

### Participants

Based on the lower bound of a bootstrapped 95% confidence interval of a pilot data set (*N* = 23), we found that the effect size of Memory Load on Bias, Guess Rate and Precision (parameters of the extended mixture model described below) were *f* = 0.53, 0.51, and 0.31, respectively. A power analysis conducted with G*Power (Faul et al., 2007) revealed that a sample of 24 participants would be required in order for the weakest Memory Load effect to be detected with a power of 90% and an alpha level of .02. However, here we aim for highly precise measurements, and we also don’t have an a priori estimate for the size of the effect of Retention Interval; therefore, we recruited 30 participants, and a large number of trials (1,600) per participant. This large data set allows ourselves and others to conduct fine-grained post-hoc analyses of the distribution of responses.

In total, 45 participants completed the experiment, from which 15 participants were excluded based on the average rating criterion (see the criterion in *Data processing and exclusion criteria*); finally, 30 participants were included for analysis. Participants were recruited from an online participant pool (Prolific, www.prolific.co), and completed the experiment in exchange for a payment of £7.5 per hour. All participants had normal or corrected-to-normal acuity and color vision. The study was approved by the local ethics review board of the University of Groningen (PSY-2122-S-0014). Participants provided informed consent before the start of the experiment.

### Stimuli, design and procedure

Each trial started with a 500ms memory display consisting of colored circles (radius: 50 px) placed evenly in a circular arrangement (radius: 250 px) around a fixation sign (*Figure 1*). The number of memory items varied from one to four, depending on the Memory Load condition. The memory colors were randomly drawn from a HSV (hue-saturation-value) color circle with full value (i.e., brightness) and saturation for each hue. (Luminance ranged from 49 cd/m^2^ to 90 cd/m^2^ on a typical lab monitor. In the present study these values varied for each participant depending on their monitor.) Participants were instructed to remember the exact colors of the memory items. Next, following a delay of 100, 500, 1,000, or 2,000 ms depending on the Retention Interval, a probe was shown for 300 ms, pointing at a location where one of the colored circles was presented^1^. Finally, participants selected the exact shade of the cued memory color on a color circle with no time limit. The color circle was randomly rotated on each trial. Visual feedback followed, showing both the selected color and the color that was actually presented at the cued location. The only constraint to the randomization process is that we used the same randomly generated set of target hues (only for the probed item) and color-circle rotations for each combination of memory load and retention interval, so as to reduce noise by making the different conditions as comparable as possible.

**Figure 1.**
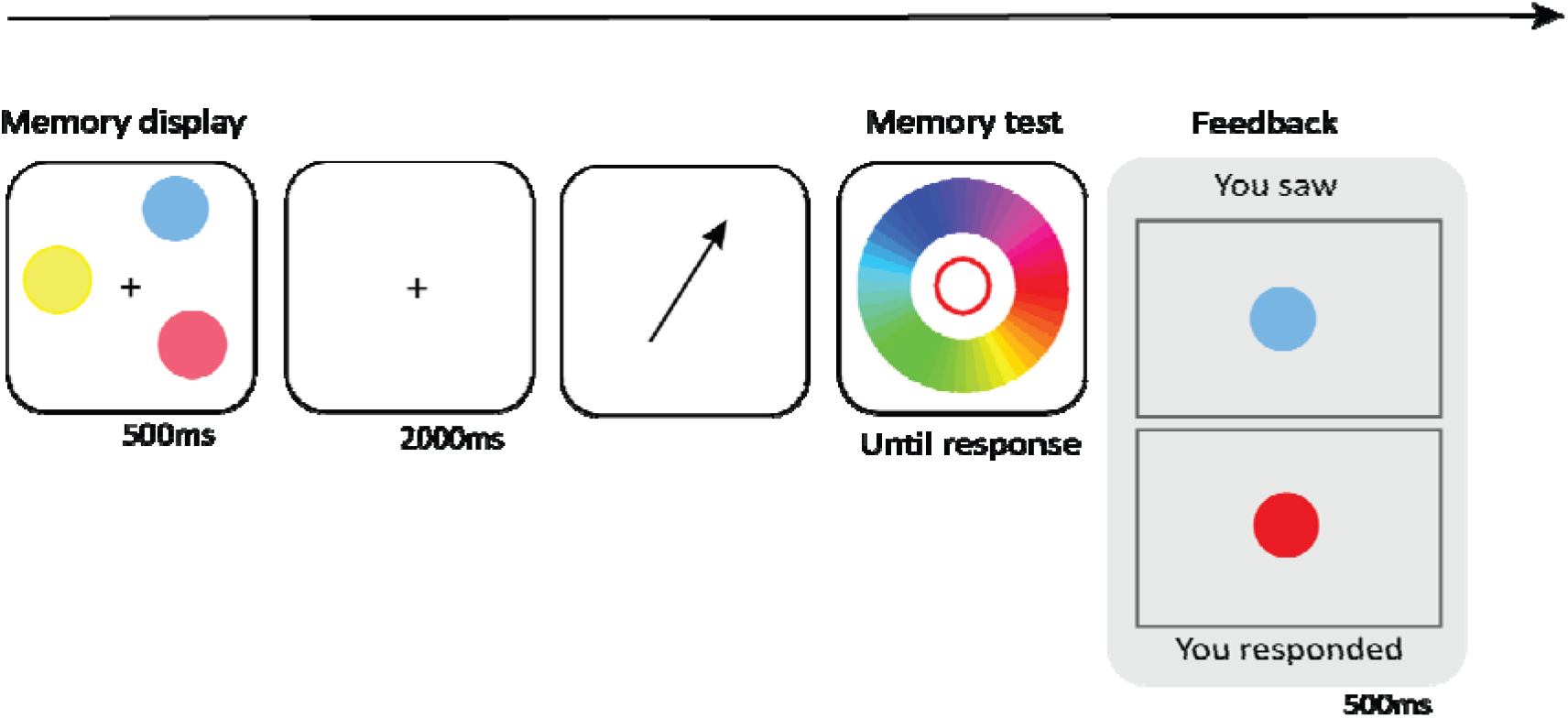
Sequence of events of a trial with a Memory Load of three and Delay Period of 2,000 ms.

Following the main experiment, we asked participants to indicate the boundaries between color categories (i.e. the hues that they perceived as boundaries between different color categories) and the prototypical color within each color category on a color wheel. Each participant indicated each boundary and prototype twice. We used the following color categories: red, pink, purple, blue, green, yellow, and orange. This data was used for our primary analyses.

Finally, we asked participants to freely partition the color wheel into as many different categories as they perceive, and then indicate the prototypical colors for each of these categories. This data was intended for an exploratory analysis, in which we explored if such unconstrained ratings are suitable for this kind of analysis.

The four levels of Memory Load and Retention Interval were mixed randomly within blocks without constraints. Participants completed four sessions in the experiment; they were free to spread these sessions across multiple days, or to run several sessions in one sitting. Within each session, there were 16 blocks of 25 trials each (400 trials each session; 1,600 trials in total); participants were free to take short breaks between blocks, but they had to finish a session in one sitting, within a maximum time window of around 60 min (the exact maximal duration is determined by Prolific). Stimulus presentation and response collection was controlled with OpenSesame / OSWeb (version 3.3; Mathôt et al., 2012).

### Data processing and exclusion criteria

For each participant, we tested if the Absolute Error (i.e. the distance between the response hue and the target hue) was significantly below chance level (lower absolute error is better). Specifically, we shuffled the response hue value of all trials, and then determined the “shuffled absolute error” based on the distance between the shuffled response hue value and the target hue value on each trial. Next, we tested if the response error of each participant was significantly lower than the shuffled response error with an independent sample t-test and an alpha level of .05. No participants were excluded based on this preregistered criterion of having an absolute error that is not below chance level.

If any of the mean color-category boundaries or prototypical colors as rated by a participant deviates more than 2 SD from the grand mean ratings, that participant was excluded. Excluded participants were replaced. Fifteen participants were excluded based on this criterion.

Individual trials were excluded when the actual duration of the delay period (as logged by the browser) deviated more than 50 ms from the intended duration. Only nine trials were excluded based on this criterion. This exclusion criterion was not pre-registered, but we considered it necessary to add after noticing that on a small proportion of trials there were temporal inaccuracies that were sufficiently large to warrant exclusion.

At first, the instructions of the color-categorization task (in which participants indicated the color boundaries and prototypical colors) were sometimes shown outside the visible area of the browser window. Therefore, we updated the task to solve this problem and asked those participants who had already participated to perform the color-categorization task again. For subsequent participants, we used the updated color-categorization task.

We further determined the Response Bias for each trial. This is identical to the Absolute Error, except that it has a sign such that positive values reflect an error in the direction of the prototypical hue of the color category to which the target hue belongs (whereas the Absolute Error is always positive). For example, say that the target hue is an orange-like shade of red, then a positive Response Bias would indicate that the response hue was shifted in the direction of a prototypical shade of red. For this analysis, we used the individually determined category boundaries and prototypes to take into account individual differences and the fact that colors are rendered slightly differently on different monitors.

### Biased Memory Model

We used a mixture model consisting of a von Mises (similar to a normal distribution for rotational values) and a uniform distribution to analyze the results. The model includes three parameters: Guess Rate ([0, 1]), such that a value of 0 indicates a pure von Mises distribution, a value of 1 indicates a pure uniform distribution, and values in-between indicate a mixture of both; Bias ([-180, 180]), which corresponds to the mean of the von Mises distribution, such that positive values indicate a shift in the direction of prototypical colors; and Precision (or kappa; κ, [0, 10,000]), which is inversely related to the variance of the von Mises distribution. For each participant, memory load, and retention interval separately, the three parameters are fitted to the Response Bias using maximum likelihood estimation. The crucial departure from our use of the mixture model, as compared to its standard use, is that we use the Response Bias, as opposed to the Error, as the dependent measure. That is, in the original model, the sign of the Error reflects the direction of the error on the color circle (clockwise or counterclockwise), which is generally not a factor of interest; in contrast, in our model, the sign of the Response Bias reflects the direction of the error relative to the corresponding prototypical color. This provides us with a meaningful and easily interpretable Bias parameter that reflects the extent to which VWM is biased by color categories. This model is freely available as a Python implementation through https://github.com/smathot/biased_memory_toolbox/.

The classic mixture model has been extended in other ways by various other researchers. Notably, Bays, Catalao, and Husain (2009) adapted the model to account for “swap errors” in which participants report a color that was memorized but not cued as the to-be-reported target. We implemented the Bias parameter also in this extended model (which is therefore able to capture both categorical biases and swap errors), and used this for exploratory analyses to test if the Bias parameter again improves the model (using an alpha level of .02). However, because of the complexity of this doubly extended mixture model, we will not make this the focus of our primary analyses.

### Statistical Analysis and Predictions

We conducted three separate repeated measure analyses of variance (RM-ANOVA) with Memory Load and Retention Interval as independent variables, and each of the three model parameters as dependent variables: Guess rate, Bias, and Precision. We used an alpha level of .02. We predicted a traditional effect of Memory Load and Retention Interval on Guess Rate and Precision, such that Guess Rate increases with Memory Load and Retention Interval while Precision decreases (in other words, it is more difficult to remember more items). Crucially, we also predicted an effect of Memory Load and Retention Interval on Bias, such that Bias increases with memory load and retention interval. This reflects our key prediction that VWM relies progressively more on categorical representations as memory load and/ or retention interval increases. We did not have specific predictions about the interaction between Memory Load and Retention Interval.

In addition, we compared the extended mixture model (with the Bias parameter) to the original mixture model (without the Bias parameter), using Akaike’s Information Criterion (AIC). We predicted that the AIC favors the inclusion of the Bias parameter. This model comparison is important, because it allows us to confirm that the Bias parameter really adds explanatory power to the model, rather than that it ‘steals’ explanatory power from one of the other parameters.

Finally, as a supplementary analysis that does not rely on the assumptions behind the mixture model, we conducted linear mixed effects models (LMER) using the R package lmerTest with Response Bias as dependent variable (determined as described above), Memory Load (reference level: 1) and Retention Interval (reference level: 100 ms) as continuous fixed effects, with random by-participant intercepts. We used an alpha level of .02.

#### Results and discussion

First, to test whether there was reliable categorical bias overall, we conducted one-sample t-tests against 0 for the Bias parameter, separately for each combination of Memory Load and Retention Interval. This showed a positive bias in all cases (all *p* < .001).

Next, we investigated how Bias was modulated by Memory Load and Retention Interval. We found an effect of Memory Load on Bias (*F*(3, 87) = 7.17, *p* < .001), such that on average Bias increased with memory load. However, as shown in *Figure 2a*, we found that the effect of Memory Load on bias was dependent on Retention Interval (Memory Load x Retention Interval: *F*(9, 261) = 2.61, *p* = .007); specifically, at the shorter intervals, we observed no significant difference in Bias as a function of Memory Load (simple main-effects analyses: 100 ms: *F*(3,87) = 1.42, *p* = .24; 500 ms: *F*(3,87) = 1.93, *p* = .13); however, at the longer intervals, bias did depend on Memory Load (1,000 ms: *F*(3,87) = 5.83, *p* = .001; 2,000 ms: *F*(3,87) = 7.00, *p* = < .001). There was no evidence for a main effect of Retention Interval on Bias (*F*(3,87) = 0.10), *p* = .96).

**Figure 2.**
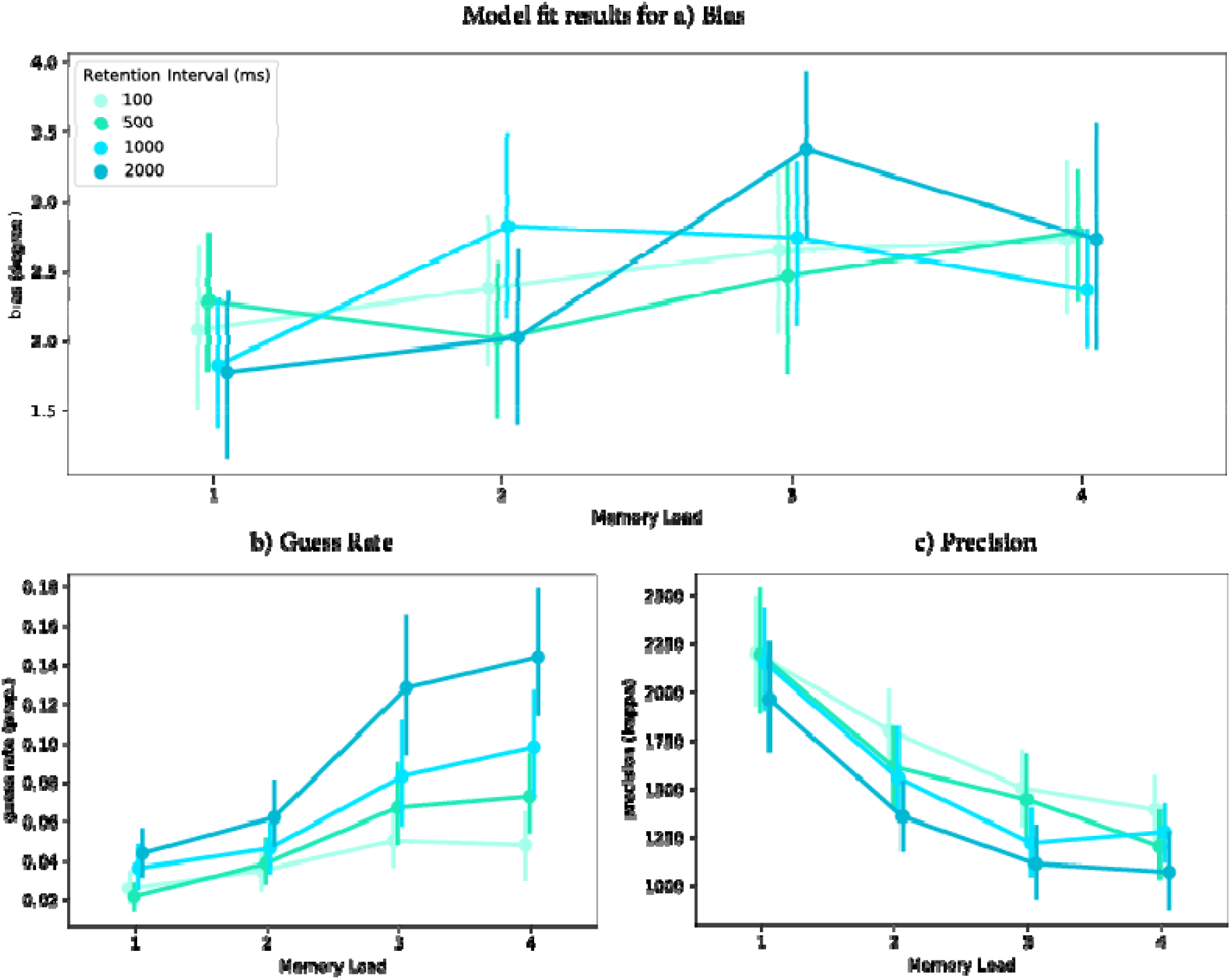
Model fit results for a) Bias, b) Guess Rate, and c) Precision as a function of Memory Load and Retention Interval. Error bars indicate bootstrapped 95% within-subject confidence intervals.

For Guess Rate, there was an effect of both Memory Load (*F*(3, 87) = 34.92, *p* < .001) and Retention Interval (*F*(3, 87) = 29.20, *p* < .001), such that guess rate increased with memory load (which plateaued at memory load 3) and retention interval. We also found a Memory Load × Retention Interval interaction (*F*(9, 261) = 6.37, *p* < .001); specifically, the dependence of Guess Rate on Memory Load increased with increasing Retention Interval.

For Precision, we found an effect of both Memory Load (*F*(3, 87) = 60.35, *p* < .001) and Retention Interval (*F*(3, 87) = 20.33, *p* < .001), such that precision decreased with increasing memory load (which plateaued at memory load 3) and retention interval. There was no interaction effect between Memory Load and Retention Interval (*F*(9, 261) = 1.02, *p* = .43). The results of Guess Rate and Precision replicated well-established effects of memory load and retention interval (Bae et al., 2015; Hardman et al., 2017; Panichello et al., 2019; Pratte et al., 2017).

Overall, our results showed that VWM contents were represented more categorically as memory load increased. Interestingly, while the effect of memory load on categorical bias was weak at shorter intervals, it became stronger at longer intervals; however, the nature of the interaction was complex, and is perhaps best described by saying that categorical bias generally increases with memory load, but is strongest for intermediate-to-high memory loads (as opposed to the highest memory load) at long intervals. (We speculate on why this is the case in the General Discussion.) Moreover, we found that guess rate increased with memory load and retention interval; on the contrary, precision decreased with memory load and retention interval.

### General Discussion

The present study investigated the extent to which VWM representations of color are biased by color categories as a function of memory load and retention interval. To test this, we added *categorical bias*, a parameter that represents the extent to which VWM is biased towards category prototypes, as an additional parameter to an extension of the mixture model of VWM (Zhang and Luck, 2008). In a delayed estimation task, participants memorized one to four colors; next, after a delay of 100, 500, 1,000, or 2,000 ms, a probe indicated the location of one of the memory colors that participants needed to report; finally, participants reproduced the target color on a color wheel. We measured their response bias, which reflects the response error in the direction of the prototypical color of the category to which the memory color belonged, using color categories that were individually established for each participant. Next, we fitted the three parameters of our Biased Memory Model (Bias, Guess Rate, and Precision) to the response bias, for each memory load and retention interval separately.

As predicted, VWM overall became more biased towards category prototypes as memory load increased. Strikingly, while the effect of memory load on categorical bias was weak at shorter intervals, it became stronger at longer intervals; however, there was a complex interaction between memory load and retention interval, such that the strongest categorical bias was not observed for the highest memory load for the longest intervals, but rather for high-to-intermediate memory loads for the longest intervals. This pattern—while complex—is consistent with previous findings that continuous and categorical representations each have a limited capacity, and that this limit is different for each type of representation. Specifically, Hardman and colleagues (2017) suggested that only a single continuous representation and two categorical representations can be maintained at the same time (see also Zhou et al., 2021 for converging evidence). In the current study, it is likely that at very short retention intervals (100 and 500 ms), visual information was still stored largely in iconic memory (Sperling 1960; or perhaps in ‘fragile working memory’; Sligte et al., 2008), which has a high capacity and relies on early visual cortex (Teeuwen et al., 2021); at this point, there was little (but some) categorical bias. After longer retention intervals (1,000 and 2,000 ms), visual information was likely transferred to VWM (Bradley & Pearson, 2012), which was has been suggested (as mentioned above) to have a capacity limit of one continuous representation and two categorical representations; at this point, categorical bias became more pronounced, especially for higher memory loads. However, there was an overall drop in bias for the highest memory loads at the longest retention intervals; one possible explanation is that when the task became too difficult, information could not be held in VWM in any form (as reflected also by high guess rates), and therefore categorical bias could no longer be meaningfully determined.

Recent neural-computational models assume that VWM representations are biased through attractor dynamics, such that memories of specific locations are systematically “attracted” to nearby attractor locations (Almeida et al., 2015; Panichello et al., 2019; Wimmer et al., 2014). Here, locations can refer to actual locations in space, but also to points in feature space, such as hues in color space, as in the present study. In these models, memory errors are characterized as a mix of diffusion of random noise and systematic drift towards stable representations, or “attractors” (Panichello et al., 2019). These attractors compress VWM representations by discretizing continuous feature space into a much smaller space of discrete attractor points so that visual information can be accurately maintained in a noisy, capacity-limited system (Koyluoglu et al., 2017; Nassar et al., 2017). In the current study, this implies that continuous VWM representations were transformed into discrete categorical representations, so that these colors could be accurately represented in VWM. Notably, our results suggest that the strength of this attracting process increases with memory load and retention interval (Panichello et al., 2019). In part this is likely because, at higher memory loads, memory is more susceptible to noise, and thus results in increased random diffusion (Yu et al., 2020); at the same time, systematic drift also increases to compensate for this increased noise. As a result, the increased drift compresses VWM representations even more strongly, making them more robust to noise, but also undermining the fidelity of these representations. In the current study, this means that VWM representations of the colors were increasingly “pulled” toward the categorical prototypes as memory load increased; when the retention interval increased, the power of this attraction toward the prototypes was strengthened even further, because attraction is a process that unfolds over time; at the same time, random diffusion also increased with increasing memory load and retention interval. As a result, VMW was represented more categorically (up to a certain point, likely because at that point the task became too hard), but also less precisely and with a higher guess rate, as memory load and retention intervals increased.

Previous studies found that categorical bias occurred as early as the perceptual period, that is, even without any retention interval between the presentation of a stimulus and its reproduction (Bae, 2021; Bays, 2015). This led to the proposal that categorical bias is caused by perception of color categories, rather than actual encoding of the colors. Our results show that while VWM representations can already be biased towards categorical prototypes during early stages of perception (i.e. with very short retention intervals), this bias becomes stronger as a result of higher-order cognitive processes; possibly, categories may initially arise through biases in perception, but as soon they have been consolidated in long-term memory, in turn influence visual processing in a top-down manner.

Taken together, the present study added categorical bias as a crucial parameter to the influential mixture model of VWM; this parameter accounts for the extent to which VWM is biased towards category prototypes. Our results showed a strong overall categorical bias in VWM (Bae 2021; Bae and Luck 2019; Hardman et al. 2017; Panichello et al. 2019); crucially, we found that VWM relies increasingly on categorical representations with increasing memory load; this effect of memory load is stronger for longer retention intervals, yet it peaked only at intermediate memory loads, rather than at the highest memory load, possibly because VWM failed to maintain this information when the task became too difficult at the highest load (as reflected also by high guess rates).

## Supplementary results

### Model comparison (AIC)

For each participant separately, we determined the AIC for a mixture model with (i.e. our Biased Memory Model) and without (i.e. the traditional mixture model) the Bias parameter. This showed that AIC favors the inclusion of the Bias parameter (paired-samples t-test: *t*(29) = -6.26, *p* < .001).

### Linear mixed effects models (LMER)

LMER analyses revealed strong evidence for the effect of Memory Load on Bias (*t*(449) = 4.56, *p* = 6.55 × 10^−6^), such that bias increased with memory load. There was no evidence for the effect of Retention Interval on Bias (*t*(449) = 0.26, *p* = .79), or a Memory Load × Retention Interval interaction (*t*(447) = 1.46, *p* = .14).

### Mixture model with Swap Errors

We also fitted a ‘doubly extended mixture model’ that included both Bias and Swap Rate as additional parameters. We analyzed the results of this model in the same way as described above for our Bias Memory Model with the only exception that we had an additional Swap parameter.

There was an effect of Memory Load on Swap errors (*F*(2, 58) = 11.67, *p* < .001), such that swap error increased with memory load. There was no evidence for an effect of retention interval (*F*(3, 87) = 2.06, *p* = .11) or a Memory Load × Retention Interval interaction (*F*(6, 174) = 2.27, *p* = .04) on Swap errors. This suggests that swap errors of non-target items added deviation of responses as memory load increased.

In this model, we did not (at our prespecified alpha level of .02) find an effect of Memory Load (*F*(2, 58) = 3.37, *p* = .04), or an effect of Retention Interval (*F*(3, 87) = 0.63, *p* = .60), or a Memory Load × Retention Interval interaction (*F*(6, 174) = 3.37, *p* = .04) on Bias. There was an effect of Memory Load (*F*(2, 58) = 17.10, *p* < .001), an effect of Retention Interval (*F*(2, 58) = 25.39, *p* < .001), and a Memory Load × Retention Interval interaction (*F*(6, 174) = 4.60, *p* < .001) on Guess Rate. We also found an effect of both Memory Load (*F*(2, 58) = 37.06, *p* < .001) and Retention Interval (*F*(2, 58) = 22.82, *p* < .001) on Precision. We did not find any Memory Load × Retention Interval interaction on Precision (*F*(6, 174) = 0.82, *p* = .55).

### Analysis with free-partitioned category ratings

We did not use the free-partitioned category ratings for any additional analysis, because some participants’ free ratings were inconsistent (for example, two category prototypes were next to each other, without any boundaries in between https://osf.io/htz6m/), or highly inconsistent between two repetitions of the same participant.

### Open practices statement

All experimental data and materials can be found on the OSF (Open Science Framework): https://osf.io/jfmw3/. All experimental data and materials for the pilot data that was used in Stage 1 review can be found on the OSF: https://osf.io/ydut7/. The Stage 1 manuscript of this study can be found on the OSF: https://osf.io/vgwb5/. The Python implementation of the mixture model toolbox can be found on GitHub: https://github.com/smathot/biased_memory_toolbox/.

**Table 1.**
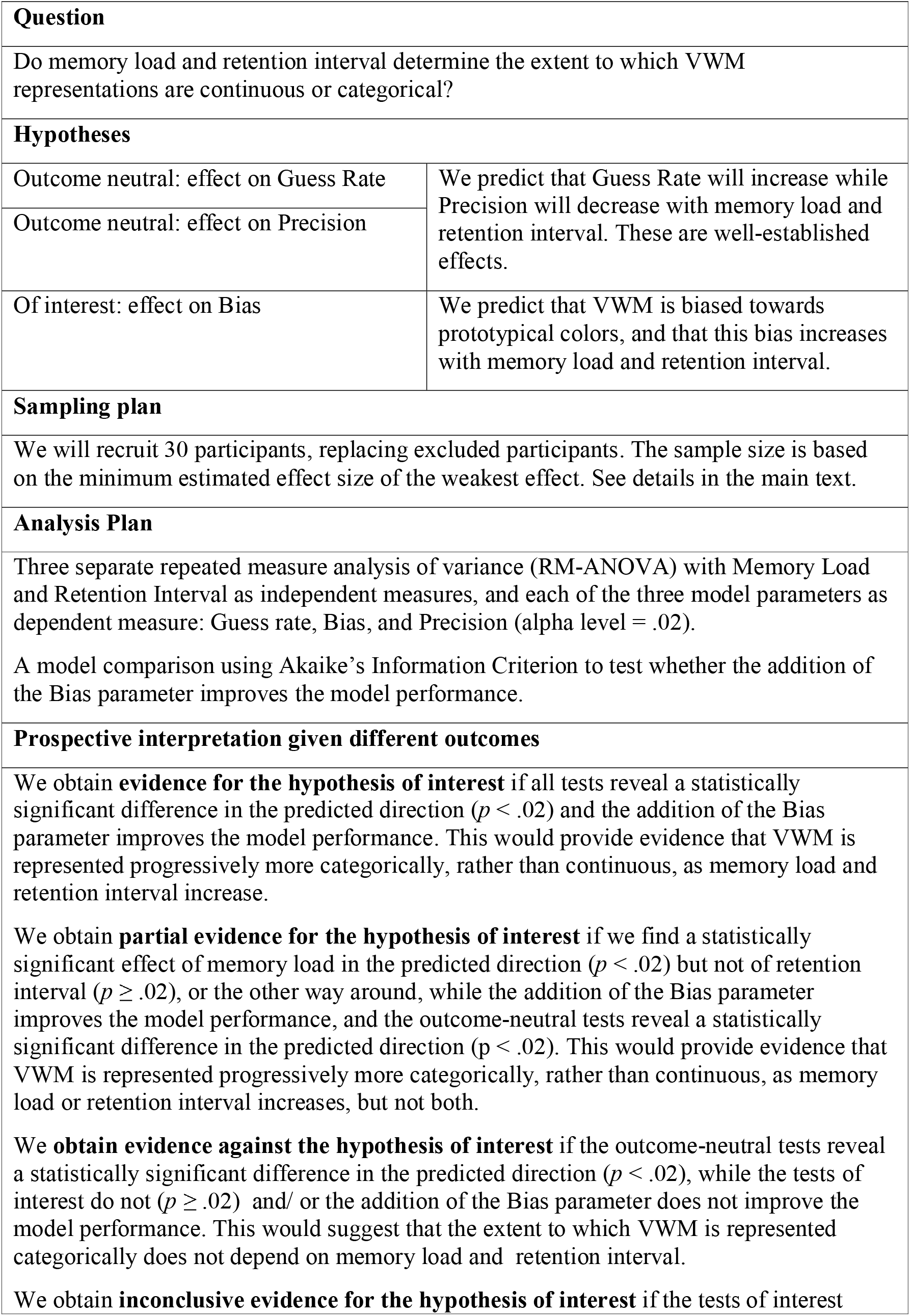
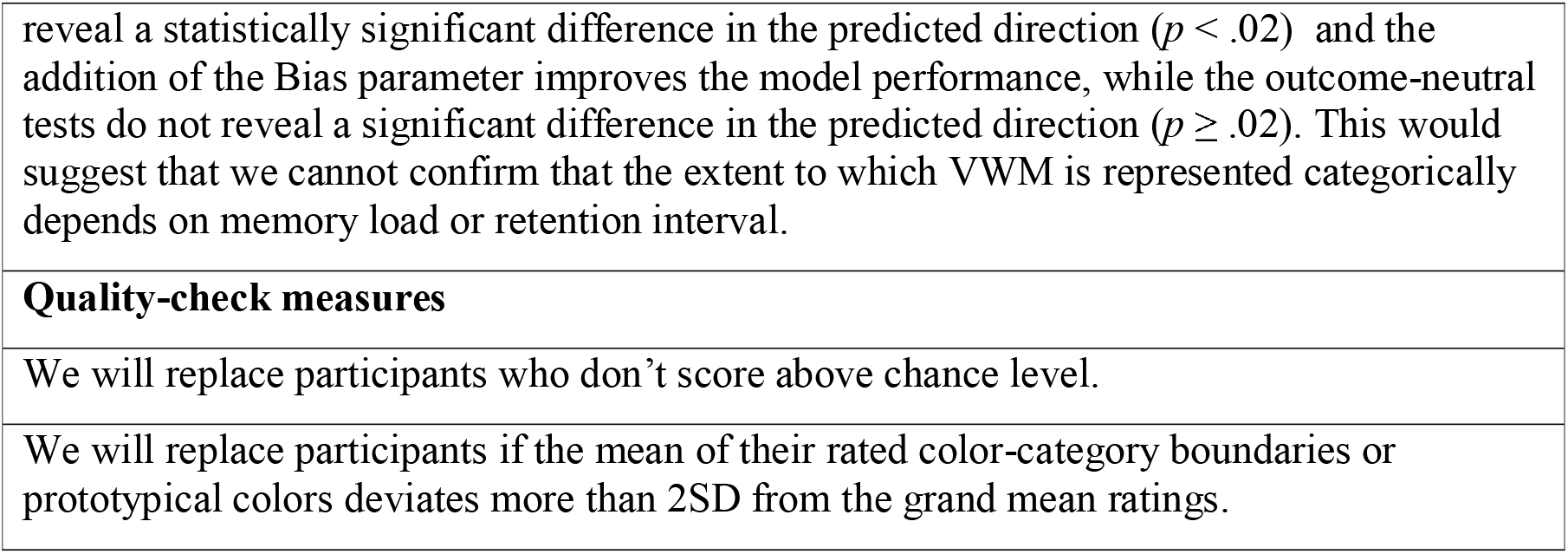
Study Design Template

## Footnotes

1 When there would be three or four potential target colors that the participant can report, it would be difficult to detect whether response errors come from bias or if the participants select randomly on the color circle, simply because any response is likely to be near a target color. Therefore, we choose to use a probe to minimize this issue by having only one target color on each trial.

